# Single olfactory receptors set odor detection thresholds

**DOI:** 10.1101/341099

**Authors:** Adam Dewan, Annika Cichy, Jingji Zhang, Kayla Miguel, Paul Feinstein, Dmitry Rinberg, Thomas Bozza

## Abstract

In many species, survival depends on olfaction, yet the mechanisms that underlie olfactory sensitivity are not well understood. Here, we examine how a conserved subset of olfactory receptors, the trace amine-associated receptors (TAARs) determine odor detection thresholds of mice to amines. We find that deleting all TAARs, or even single TAARs, results in significant odor detection deficits. This finding is not limited to TAARs, as the deletion of a canonical odorant receptor reduced behavioral sensitivity to its preferred ligand. Remarkably, behavioral threshold is set solely by the most sensitive receptor, with no contribution from other highly sensitive receptors. In addition, increasing the number of sensory neurons (and glomeruli) expressing a threshold-determining TAAR does not improve detection, indicating that sensitivity is not limited by the typical complement of sensory neurons. Our findings demonstrate that olfactory thresholds are set by the single highest affinity receptor, and suggest that TAARs are evolutionarily conserved because they determine the sensitivity to a class of biologically relevant chemicals.

## Introduction

No artificial chemical detector can match the simultaneous breadth, flexibility and sensitivity of biological olfactory systems^1^. In terrestrial vertebrates, chemicals in the environment (odorants) are inhaled into the nasal cavity where they are recognized by olfactory sensory neurons (OSNs) in the olfactory epithelium^2^. In mice, each OSN expresses one receptor from a large repertoire of olfactory receptor genes^3^, and OSNs that express the same receptor send convergent axonal projections to defined glomeruli in the olfactory bulb^4^.

It is currently unknown how molecular recognition at the level of receptors and OSNs gives rise to the exquisite sensitivity that animals display at the behavioral level. Previous data show that OSNs exhibit modest sensitivities, responding to odorants in the micromolar or nanomolar concentration range^5, 6^, while observed behavioral detection thresholds can be orders of magnitude lower^7, 8^. Part of this disparity may be attributed to long-standing difficulties in functionally expressing olfactory receptors and identifying the most sensitive (threshold-determining) receptors for specific odorants. While previous experiments have shown detection deficits by removing individual (or sets of) olfactory receptors^9, 10, 11^, in most species (particularly in mammals) it remains unclear what factors (e.g. receptor sensitivity, sensory neuron number, glomerular convergence) influence behavioral detection thresholds.

In mice, there are two classes of main olfactory G-protein coupled olfactory receptors: a large family of >1,000 canonical odorant receptors (ORs) and the small family of 14 trace amine associated receptors (TAARs). The TAARs are phylogenetically distinct from ORs, being more closely related to biogenic amine receptors. However, both classes of receptors couple to the same olfactory signal transduction cascade^12, 13^, and OSNs that express TAARs or ORs project axons to typical glomeruli in the main olfactory bulb^14, 15^. The TAARs respond selectively to amines—a class of compounds that is ubiquitous in biological systems—and have been implicated in mediating innate responses to odorants that act as social and predator-derived cues in mice^16, 17^. However, the TAARs are highly conserved across vertebrates^18^ and likely serve a more common critical function in olfaction.

We have recently shown that the TAARs exhibit high sensitivity, responding to amines at subpicomolar concentrations^12^, and that TAAR glomeruli are the most sensitive amine-responsive glomeruli in the dorsal olfactory bulb^14, 19^. This suggests that the TAAR family, and perhaps even single TAARs, may contribute significantly to amine sensitivity at the behavioral level.

To determine how TAARs contribute to olfactory sensitivity, we characterized the response specificity of TAAR glomeruli, and measured odor detection thresholds in mice that lack all TAARs, or single TAAR or OR genes. This approach allowed us to rank order the sensitivity of specific TAARs to defined odorants, and to systematically quantify how much each receptor contributes to sensitivity. We find that a single TAAR or canonical OR can set the behavioral detection threshold to a given odorant. Our data demonstrate for the first time, a unique contribution of TAARs to olfaction that may be common to all vertebrates.

## Results

### The TAAR family and amine detection

To measure how the TAAR gene family contributes to amine detection, we examined TAAR cluster deletion mice (ΔT2-9) which lack all 14 olfactory TAARs (Figure 1a)^14^. To measure behavioral detection thresholds rigorously, we developed a head-fixed, go/no-go operant conditioning assay combined with well-controlled and highly-reproducible stimulus delivery ^20^. Mice report the absence or presence of an odor by licking or not licking for a water reward (Figure 1b). To control for differences in task performance due to genetic background, we compared mutants to wild-type littermates of the same strain. Using this approach, we first examined the sensitivity of ΔT2-9 mice to two TAAR ligands phenylethylamine and isopentylamine, and to a non-TAAR ligand methyl valerate (Figure 1c-e). Overall, wild-type mice were highly sensitive to all odors tested, exhibiting accurate responses to concentrations below 1×10^−10^*M*, well below the sensitivity of the human experimenters. For simplicity, we compare sensitivity across animals using the odor concentration at half-maximal behavioral performance (C_½_).

**Figure 1.**
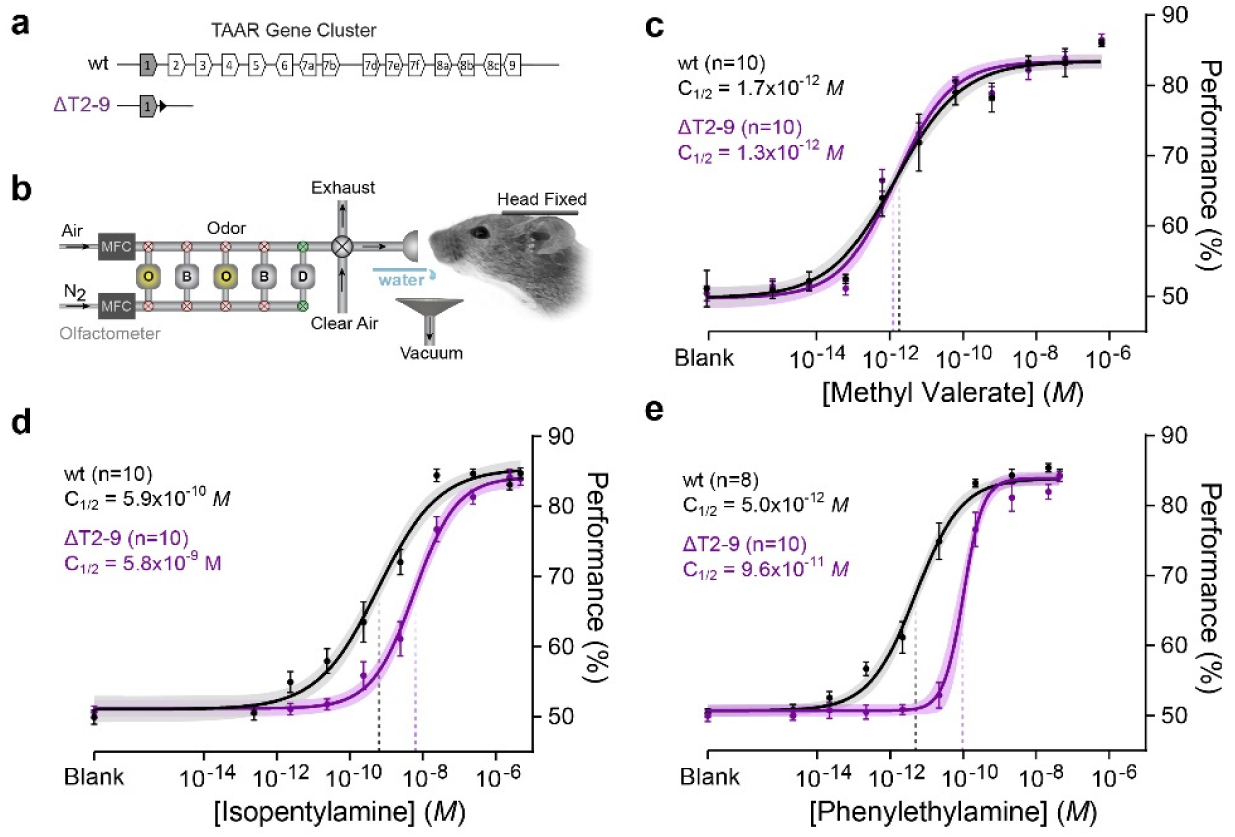
Removing all olfactory TAARs causes detection deficits to amines. **(a)** Diagram of the wild-type mouse TAAR gene cluster(wt), and cluster deletion (ΔT2-9). Olfactory TAARs (white) and non-olfactory *Taar1* (shaded) are shown (polygons reflect gene orientation). LoxP site indicated as black triangle. **(b)** Schematic of odor delivery for thresholding in head-fixed mice. Flow dilution olfactometer switches between a pressure-balanced dummy (D) carrier vial (via normally open valves, green) and either odor (O) and blank (B) vials (via normally closed valves, red). During stimulus application, a final valve re-directs pressure-balanced, odorized air from exhaust to the animal. Mice lick for water reward in go/no-go detection task. (**c-e**) Psychometric curves of wild-type (black) and mice lacking all olfactory TAARs (purple) to two amines (isopentylamine and phenylethylamine) and one non-amine (methyl valerate). Plots show mean +/- SE with a shaded 95% confidence interval. Data were fitted with a Hill function. Behavioral sensitivity is defined as the odor concentration at half-maximal behavioral performance (C_½_) during a head-fixed Go No-Go conditioned assay. Methyl valerate: *wt* C_½_ = 1.7 × 10^−12^ *M* (95% CI = 0.9-3.5×10^−12^ *M*); *ΔT2-9* C_½_= 1.3×10^−12^ *M* (95% CI = 0.7-2.2×10^−12^ *M*). Isopentylamine: *wt* C_½_= 5.9×10^−10^ *M* (95% CI = 3.0-11.2×10^−10^*M*); *ΔT2-9* C_1/2=_ 5.8×10^−9^ *M* (95% CI = 3.7-9.0×10^−9^ *M*). Phenylethylamine: *wt* C_½_= 5.0×10^−12^*M* (95% CI = 3.2-7.6×10^−12^*M*); *ΔT2-9* C_½_= 9.6×10^−11^ *M* (95% CI = 6.2-12.1×10-11*M*).

Deleting the TAARs caused an 18.4-fold increase i n C_½_ for phenylethylamine (Figure 1e): 5.0×10^−12^*M* for wild-type and 9.6×10^−11^ *M* for *ΔT2-9* mice (p=1.5×10^−12^; F=60.49, Sum-of-squares F-test). TAAR deletion also caused a 10-fold increase in C_½_ to isopentylamine (Figure 1d): 5.9×10^−10^ *M* for wild-type and 5.8×10^−9^ *M* for *ΔT2-9* mice (p=1.7×10^−8^; F=34.96, Sum-of-squares F-test). No difference in sensitivity was observed for the non-amine methyl valerate (Figure 1c): C_½_ = 1.7 × 10^−12^ *M* for wild-type vs. 1.3×10^−12^ *M* for *ΔT2-9* (p=0.37; F=0.81, sum-of-squares F-test). These data indicate that the TAAR gene family normally sets the detection thresholds of mice to phenylethylamine and isopentylamine.

### Odorant Response Profiles of TAARs in awake mice

To characterize individual TAAR genes that are likely critical for detecting specific amines, we focused on TAAR3, TAAR4 and TAAR5—receptors that are highly sensitive to isopentylamine, phenylethylamine, and trimethylamine, respectively^12, 13, 14, 19^. However, all previous data on TAAR specificity have come from *in vitro* studies, or from *in vivo* glomerular imaging in anesthetized mice.

To characterize TAAR response profiles under behaviorally relevant conditions, we measured odor-evoked responses from genetically-identified glomeruli in the dorsal olfactory bulb using calcium imaging in awake mice (Figure 2a, b). Ten out of the 14 olfactory TAARs (including TAAR 3, 4, and 5) have corresponding dorsal glomeruli within our imaging window, while TAAR 6, 7a, 7b and 7d likely have glomeruli outside the window^14, 15, 21^. We imaged a strain of mice in which all glomeruli express the calcium indicator GCaMP3 and in which TAAR3 and TAAR4 glomeruli are differentially labeled with YFP and RFP, respectively (Figure 2a). In the same mice, TAAR5 glome ruli were identified based on data from separate GCaMP imaging experiments in which TAAR5 glomeruli were labeled with RFP. It should be noted that linkage of these clustered genes precludes crossing three tagged alleles into the same mouse.

**Figure 2.**
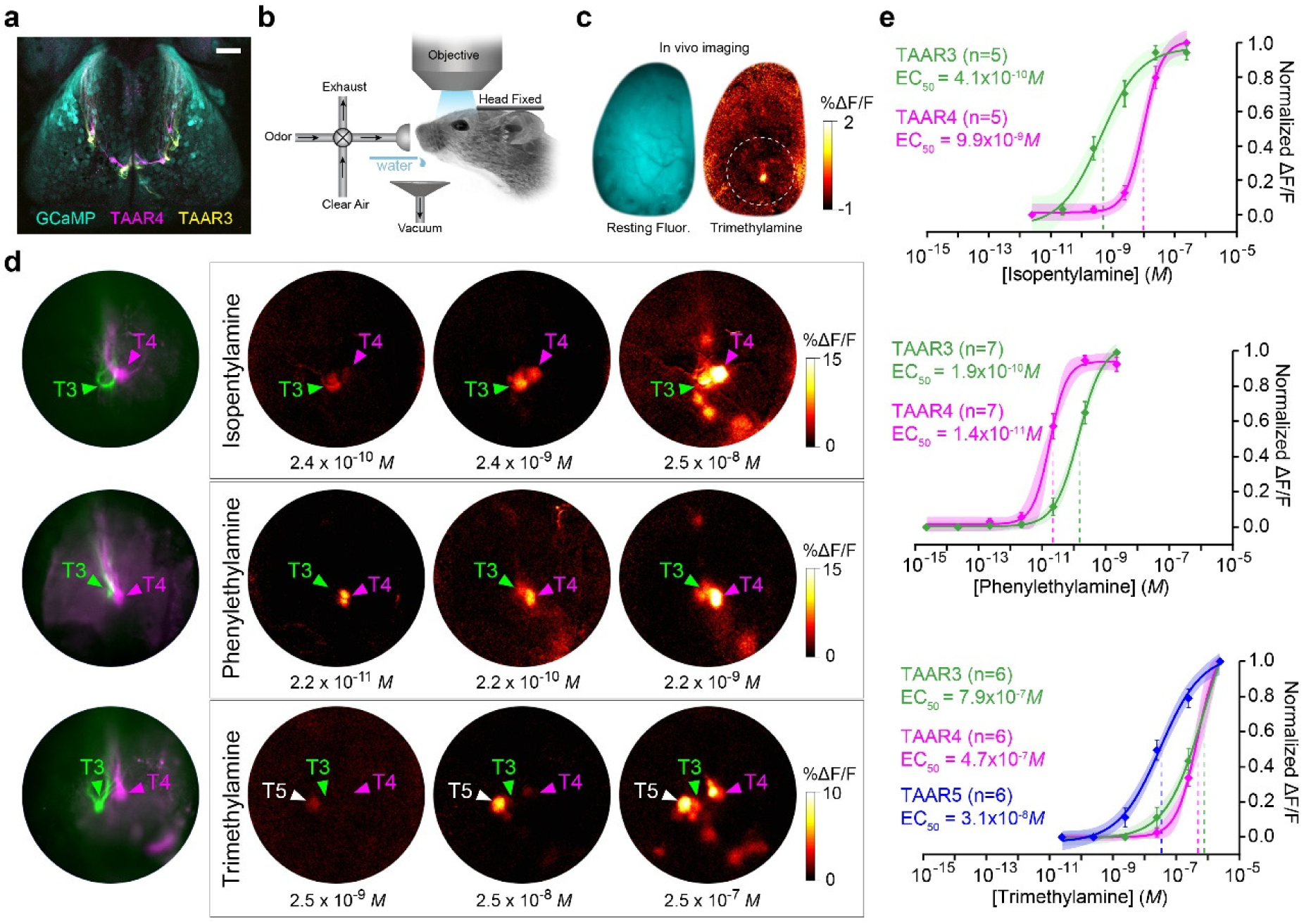
TAAR glomeruli are the most sensitive amine receptors in the dorsal olfactory bulb. **(a)** Confocal image of the dorsal olfactory bulbs from a compound heterozygous *OMP-GCaMP3/TAAR3-YFP/TAAR4-RFP* mouse. Anterior is up. **(b)** Schematic of setup for wide-field *in vivo* calcium imaging in head-fixed, awake mice. Odors are delivered using a flow dilution olfactometer, and mice receive water rewards after odor trials. **(c)** Wide-field fluorescence image of an olfactory bulb showing resting GCaMP fluorescence (cyan) and response in a single glomerulus to trimethylamine (warm colors) in an awake mouse. Circle shows the general region of the olfactory bulb imaged at higher magnification in all experiments. (**d**) Higher magnification imaging of odor-evoked activity from TAAR glomeruli in the dorsal olfactory bulbs. (left) Resting fluorescence shows genetically tagged TAAR3 (green) and TAAR4 (magenta) glomeruli in three different mice. (right) Odor evoked GCaMP responses to amines. Medial is to the right. Locations of tagged glomeruli are marked with colored arrows. Locations of TAAR5 glomeruli, based on response profile, are marked with white arrows. **(e)** Dose-response curves from individual TAAR glomeruli to isopentylamine (top), phenylethylamine (middle), and trimethylamine (bottom). Plots show mean +/- SE with a shaded 95% confidence interval. Data were fitted to a Hill function. Isopentylamine: TAAR3 EC_50_ = 4.1 × 10^−10^ *M* (95% CI = 1.3-9.9 × 10^−10^ *M*); TAAR4 EC_50_ = 9.9x 10^−9^ *M* (95% CI = 6.9-14 × 10^−9^ *M*). Phenylethylamine: TAAR4 EC_50_ = 1.4 × 10^−11^ *M* (95% CI = 1.5-2.4 × 10^−11^ *M*); TAAR3 EC_50_ = 1.9 × 10^−10^ *M* (95% CI = 1.0-1.8 × 10^−10^ *M*). Trimethylamine: TAAR5 EC_50_ = 3.1 × 10^−8^ *M* (95% CI 1.7-7.6 × 10^−8^ *M*); TAAR4 EC_50_ = 4.7 × 10^−7^ *M* (95% CI 1.2-21.0 × 10^−7^ *M*); TAAR3 EC_50_ = 7.9 × 10^−7^ *M* (95% CI 2.9-12.8 × 10^−7^ *M*). Scale bar in A = 500 µm in A and C, 200 µm in D.

At low concentrations, isopentylamine, phenylethylamine, and trimethylamine strongly and specifically activate TAAR3, TAAR4, and TAAR5 glomeruli, respectively without eliciting a response in other dorsal glomeruli^14, 19^ (Figure 2c). With increasing concentrations, response profiles exhibited some overlap. Both isopentylamine and phenylethylamine robustly activated both TAAR3 and TAAR4 glomeruli (Figure 2d). Across concentration, TAAR3 was 24-fold more sensitive to isopentylamine than TAAR4 (Figure 2e). The EC_50_ values were 4.1 × 10^−10^ *M* for TAAR3 vs. 9.9x 10^−9^ *M* for TAAR5 (p=1.2x 10^−6^, F=31.65, Sum-of-squares F-test). Likewise, TAAR4 was 8-fold more sensitive to phenylethylamine than TAAR3 (Figure 2e). The EC_50_ values were 1.4 × 10^−11^ *M* for TAAR4 vs. 1.9 × 10^−10^ *M* for TAAR3 (p=1.4×10^−12^, F=65.31, Sum-of-squares F-test). Thus, at low concentrations, phenylethylamine and isopentylamine differentially activate TAAR3 and TAAR4. Similarly, trimethylamine activates all three TAARs, but TAAR5 was 20- fold more sensitive to trimethylamine than either TAAR3 or TAAR4 (Figure 2e). The EC_50_ values were 3.1 × 10^−8^ *M* for TAAR5 vs. 4.7 × 10^−7^ *M* for TAAR4 (p=1.8×10^−8^, F=40.95, Sum-of-squares F-test) and 7.9 × 10^−7^ *M* for TAAR3 (p=5.7×10^−4^, F=13.15, Sum-of-squares F-test). TAAR3 and TAAR4 did not differ in their sensitivity to trimethylamine (p=0.594, F=0.29, Sum-of-squares F-test). These results define for the first time the relative sensitivities of TAAR3, TAAR4, and TAAR5 to amines in awake animals.

We were surprised to observe a general concordance between the sensitivity of the most sensitive TAAR glomeruli and the behavioral detection thresholds of the animals (within a factor of 2-5 fold) (Supplementary Fig. 1). While there are some methodological differences that might impact a detailed comparison of glomerular and behavioral sensitivity (see Methods), the general agreement between glomerular and behavioral sensitivity suggested that TAAR3, TAAR4 and TAAR5 might be responsible for setting detection thresholds for their highest affinity ligands.

### Sensitivity deficits from single TAAR deletions

To determine whether individual TAARs contribute significantly to amine detection thresholds, we examined strains of mice in which the coding sequences for TAAR3, TAAR4 or TAAR5 were replaced with coding sequences for different reporters (Figure 3a; see Methods). We observed significant decreases in behavioral sensitivity caused by deleting each of the three TAARs (Figure 3b-d). Isopentylamine preferentially activates TAAR3, and mice lacking only TAAR3 (ΔT3) exhibited a 6.3-fold decrease in sensitivity to isopentylamine (Figure 3b) with C_½_ values of 1.5 × 10^−9^ *M* for wild-type vs. 9.4 × 10^−9^ *M* for *ΔT3* (p=1.3×10^−12^, F=61.09; Sum-of-squares F-test). Similarly, phenylethylamine preferentially activates TAAR4, and mice lacking only TAAR4 (ΔT4) exhibited a 6.9- fold decrease in sensitivity to phenylethylamine (Figure 3c). C_½_ values were 5.9 × 10^−12^ *M* for wild-type vs. 4.0 × 10^−11^ *M* for *ΔT4* (p=0.005, F=8.32, Sum-of-squares F-test). In the case of trimethylamine, which preferentially activates TAAR5, we note that thresholding with this odorant was technically challenging due to its high volatility. However, we were able to measure a 4-fold reduction in sensitivity to trimethylamine in mice lacking TAAR5 (Figure 3d). C_½_ values were 2.6 × 10^−8^ *M* for wild-type vs. 8.6 × 10^−8^ *M* for *ΔT5* (p=0.028, F=4.9, Sum-of-squares F-test). These data directly demonstrate that single TA ARs contribute significantly to behavioral sensitivity.

**Figure 3.**
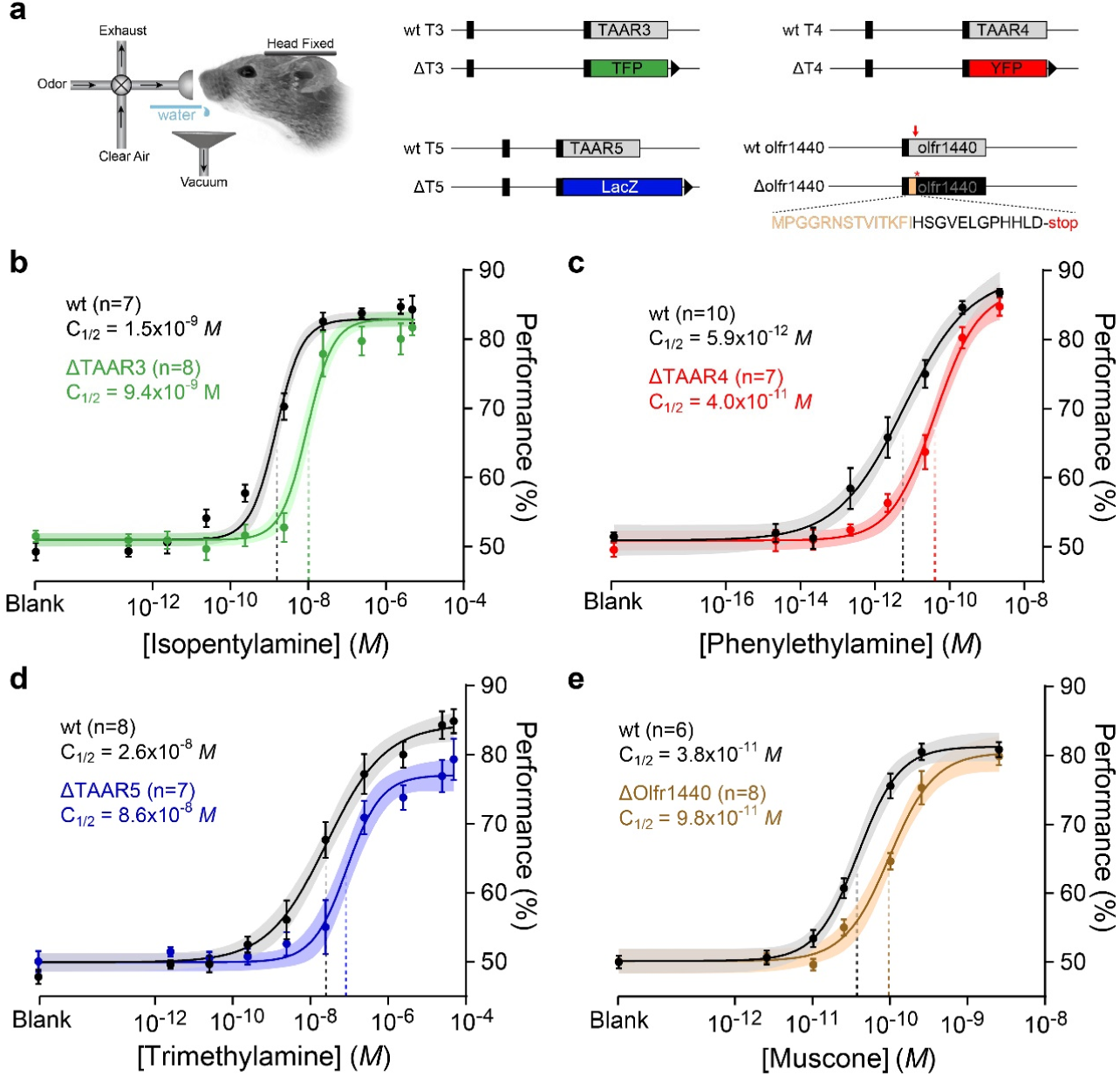
Deletion of single olfactory receptor genes causes detection deficits. (**a**) Head-fixed thresholding was performed on mice harboring different gene targeted alleles (right). Three coding sequence replacements at the *taar3*, *taar4* and *taar5* loci are shown. Black boxes are non-coding regions. Receptor coding sequences (grey) were replaced with teal fluorescent protein (TFP; green), β-galactosidease (LacZ; blue) and yellow fluorescent protein, (YFP; red). The *olfr1440* deletion was generated by CRISPR-based gene editing. Red arrow shows position of targeted PAM site and frameshift. The truncated coding sequence is shown (native sequence in tan). LoxP sites indicated by black triangles (**b-e**) Psychometric curves for receptor deletion (colored) and wild-type littermate controls (black) for ΔTAAR3 (green) to isopentylamine (**b**), ΔTAAR4 (red) to phenylethylamine (**c**), ΔTAAR5 (blue) to trimethylamine (**d**), and ΔOlfr1440 (tan) to muscone (**e**). Data were fitted using a Hill function. Behavioral sensitivity is defined as the odor concentration at half-maximal behavioral performance (C_½_) during a head-fixed Go No-Go conditioned assay. Plots show mean +/- SE with a shaded 95% confidence interval. Isopentylamine: *wt* C_½_ = 1.5 × 10^−9^ *M* (95% CI = 0.9-2.2 × 10^−9^ *M*); *ΔT3* C_½_= 9.4 × 10^−9^ *M* (95% CI = 6.6-13.2 × 10^−9^ *M*). Phenylethylamine: *wt* C_½_ = 5.9 × 10^−12^ *M* (95% CI = 2.4-18.9 × 10^−12^ *M*); *ΔT4* C_½_ = 4.0 × 10^−11^ *M* (95% CI = 2.3-9.8 × 10^−11^ *M*). Trimethylamine: *wt* C_½_ = 2.6 × 10^−8^ *M* (95% CI = 1.2-6.2 × 10^−8^ *M*); *ΔT5* C_½_ = 8.6 × 10^−8^ *M*; 95% CI 4.5-20.0 × 10^−8^ *M*). Muscone: *wt* C_½_ = 3.8 × 10^−11^ *M* (95% CI = 3.1- 5.0 × 10^−11^ *M*); *ΔOlfr1440* C_½_ = 9.8 × 10^−11^ *M* (95% CI 7.4-12.8 × 10^−11^ *M*).

### Sensitivity deficit from a single odorant receptor deletion

To compare the detection deficits from TAAR deletions with deficits from deleting canonical ORs, we characterized the effect of deleting a model class II OR, *Olfr1440* (MOR215-1). Deletion of *Olfr1440* has been reported to reduce sensitivity to muscone as measured using an investigation-based behavioral assay in freely moving mice^9^. We generated a strain of mice (*ΔOlfr1440*) by CRISPR/Cas9-mediated gene editing, in which the *Olfr1440* coding sequence is truncated due to a frameshift and premature stop codon (Figure 3a). We then measured detection thresholds in wild-type and homozygous deletion littermates. Wild-type mice exhibited an appreciable sensitivity to muscone (C_½_ = 3.8×10^−11^*M*)— similar to the behavioral sensitivity observed for amines. Moreover, homozygous *ΔOlfr1440* mice exhibited a 2.6-fold reduction in muscone sensitivity (C_½_ = 9.8 × 10^−11^ *M*; p=1.1×10^−5^, F=21.83; Sum-of-squares F-test; Figure 3e), a similar magnitude effect as seen with the single TAAR deletions. No difference in sensitivity was observed in *ΔOlfr1440* mice for the control odor methyl valerate (Supplementary Fig 2). These data indicate that the contribution of single receptors to behavioral sensitivity can be similar for TAARs and canonical ORs.

Deleting *olfr1440* has been reported to produce a much larger detection deficit than what we observe via thresholding^9^. To test whether this is attributable to the different assays, we measured the sensitivity of our ΔT2-9 mice using a similar investigation-based assay (Supplementary Fig. 3). As expected, ΔT2-9 mice displayed reduced investigation to phenylethylamine as compared with wild-types (p=0.016), but showed no difference for methyl valerate (p=0.932, Generalized Estimating Equations Model). The lowest concentration of phenylethylamine that elicited appreciable investigation was 1,000-fold higher for ΔT2-9 mice than wild-type littermates (wild-type = 1×10^−6^ *M* vs. ΔT2-9 = 1×10^−3^ *M*, Generalized Estimating Equations Model). Thus, the investigation assay reports a change in olfactory function, but appears to overestimate the size of the detection deficit (see Discussion).

### Amine sensitivity is set solely by the most sensitive TAAR

Behavioral detection thresholds could be set exclusively by the most sensitive receptor population, or might require combining (or pooling) information from several of the most sensitive receptor populations ^22^. Thus, we sought to determine whether, in the presence of the most sensitive receptor, selective removal of other high-affinity receptors impairs detection of a given odorant. We reasoned that we could address this question using the TAARs and phenylethylamine.

The TAAR cluster deletion had a significantly larger effect on phenylethylamine sensitivity than the TAAR4 deletion alone (p=3.4×10^−5^, F=18.62, Sum-of-squares F-test; wild-type comparison, p=0.870, F=0.03, sum of squares F-test; Supplementary Fig. 1). The more pronounc ed deficit in the cluster deletion indicates that phenylethylamine sensitivity in TAAR4 deletion animals is likely set by other (remaining) high-affinity TAARs. We wanted to unequivocally identify these phenylethylamine-responsive TAARs to study their effects on threshold.

Our imaging data implicate TAAR3 as the secondmost-sensitive phenylethylamine receptor in the dorsal bulb—second only to TAAR4. However, several TAARs absent in the cluster deletion have ventral glomeruli that are inaccessible for optical imaging. Therefore, we used the DREAM assay^23^ to identify whether any of these ventral TAARs respond to phenylethylamine (Figure 4a). The DREAM method identifies receptor-ligand pairs *in vivo* by measuring odor-evoked reduction in the expression of activated receptor genes.

**Figure 4.**
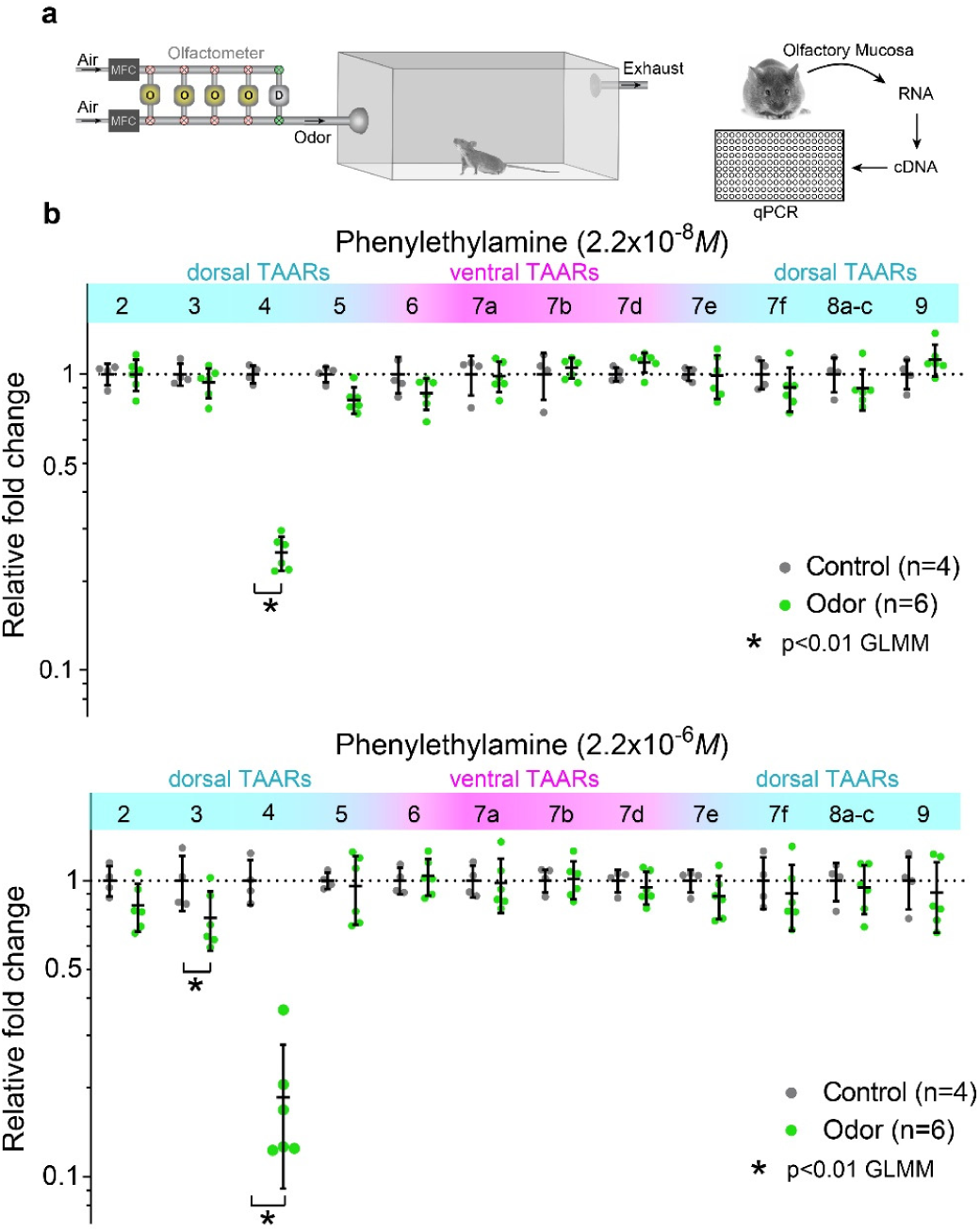
Identification of TAARs that respond to phenylethylamine using DREAM. (**a**) Diagram of experimental workflow including odor exposure, isolation of olfactory mucosa, generation of cDNA, and measurement of receptor expression via qPCR. (**b**) Plots show relative levels of TAAR gene expression in the olfactory mucosa from odorized (green) and non-odorized (control; black) mice as determined by qPCR (normalized to the average non-odorized value). Mice were exposed to phenylethylamine at two concentrations for 24 hours. Concentrations given are calculated values at the olfactometer outlet at the start of the experiment. Responsive TAARs are indicated by decreases in expression. Black lines displaying mean +/- SD are plotted over individual data points. Ventrally-expressed TAAR genes (correspond to glomeruli outside the imaged area of the dorsal bulb) are shaded in pink. Asterisk denotes p<0.01 General Linear Mixed Model.

At the lower of two concentrations tested, phenylethylamine elicited a significant reduction in TAAR4 expression only (Figure 4b). At the higher of the tested concentrations, phenylethylamine elicited a reduction in both TAAR3 and TAAR4 expression (Figure 4c). No activation was seen for any of the other TAARs. We note that both concentrations delivered to the odorization chamber were well above threshold, consistent with the idea that the DREAM effect may be related to adaptation^23^. Together with our imaging assay, the data demonstrate that TAAR3 is the second-most-sensitive (and TAAR4, the most sensitive) TAAR for phenylethylamine.

We then asked whether TAAR3 (the second most sensitive TAAR) contributes to phenylethylamine sensitivity when TAAR4 (the most sensitive TAAR) is intact. We observed that deletion of TAAR3 alone produced no change in sensitivity to phenylethylamine when TAAR4 was intact (Figure 5a): C_½_ = 3.9 × 10^−12^ *M* for wild-type vs. 3.7 × 10^−12^ *M* for *ΔT3* (p=0.923, F=0.009, sum-of-squares F-test). Thus, behavioral sensitivity to phenylethylamine is determined solely by TAAR4.

**Figure 5.**
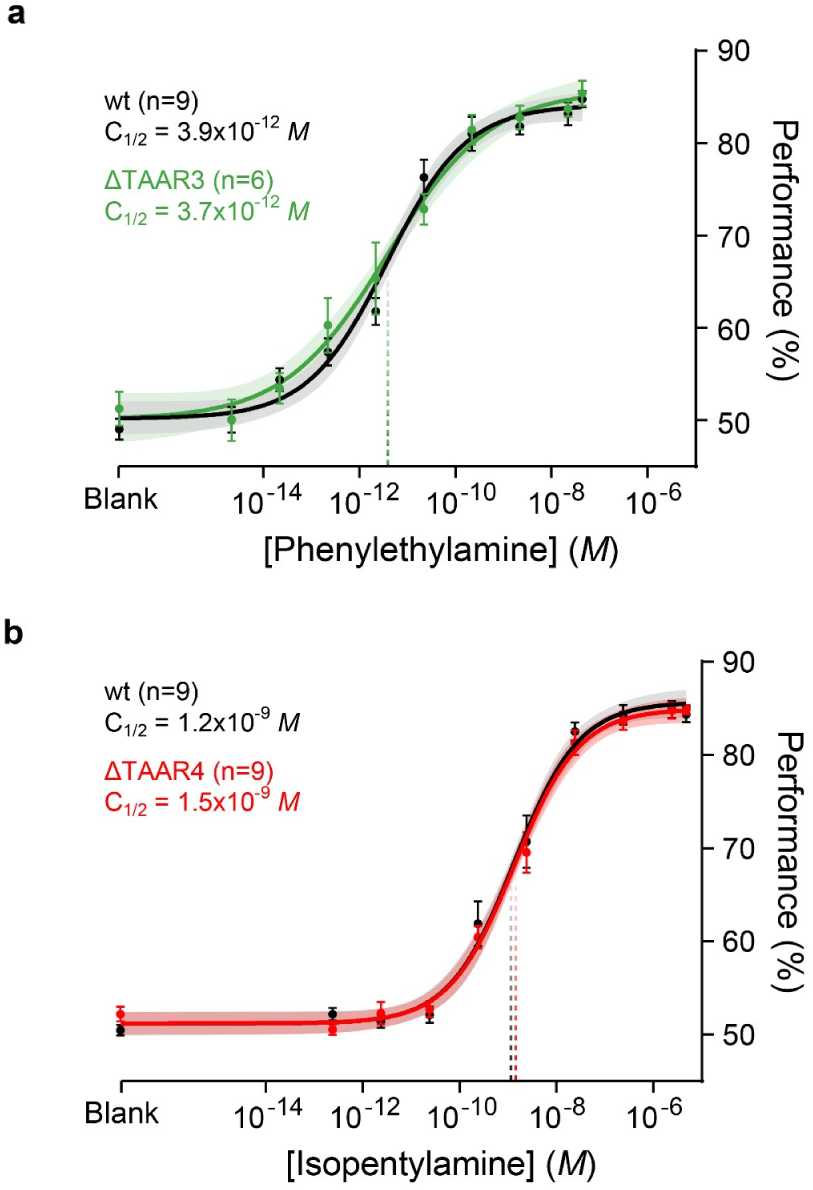
Deletion of single TAARs does not impact sensitivity to non-preferred ligands. (**a**) Psychometric curves of wild-type (black) and TAAR4 deletion littermates (red) to isopentylamine. (**b**) Psychometric curves of wild-type (black) and TAAR3 deletion littermates (green) to phenyethylamine. Plots show mean +/- SE with a shaded 95% confidence interval. Data were fitted using a Hill function. Behavioral sensitivity is defined as the odor concentration at half-maximal behavioral performance (C_½_) during a head-fixed Go No-Go conditioned assay. Phenylethylamine: *wt* C_½_ = 3.9 × 10^−12^ *M* (95% CI = 2.0-7.1 × 10^−12^); *ΔT3* C_½_ = 3.7 × 10^−12^ *M* (95% CI = 1.1-12.5 × 10^−12^). Isopentylamine: *wt* C_½_ = 1.2 × 10^−9^ *M* (95% CI = 0.6-1.9 × 10^−9^ *M*); *ΔT4* C_½_ = 1.5 × 10^−9^ *M* (95% CI 0.9-2.3 × 10^−9^ *M*).

We note that similar reasoning could not be applied for the odorant isopentylamine since the detection deficits caused by the cluster deletion and the TAAR3 deletion alone were statistically indistinguishable after compensating for differences between the strains (p=0.351, F=1.00, Sum-of-squares F-test; Supplementary Fig. 1). That is to say, we have no evidence that any of the remaining TAARs contribute to isopentylamine sensitivity when TAAR3 is deleted. Despite this, our imaging data show that TAAR4 has an appreciable affinity for isopentylamine. We therefore tested whether TAAR4 deletion alone (in the presence of TAAR3) affects isopentylamine sensitivity, and observed no effect (Figure 5b): C_½_ = 1.2 × 10^−9^ *M* for wild-type vs. 1.5 × 10^−9^ *M* for *ΔT4* (p=0.398, F=0.72; Sum-of-squares F-test). In summary, our data are consistent with the idea that phenylethylamine sensitivity is determined solely by the highest affinity TAAR (TAAR4).

### Overexpressing TAAR4 does not affect sensitivity

Our data indicate that behavioral sensitivity can be set by a single input to the olfactory system, as defined by a specific receptor and its associated OSNs and glomeruli. Previous data indicate that increasing the number of OSNs that respond to a given odorant improves sensitivity^24^. Therefore, we asked whether increasing the number of OSNs and glomeruli for the threshold-determining receptor TAAR4 can enhance behavioral sensitivity. We generated transgenic mice in which the number of OSNs that express TAAR4 is significantly increased by driving expression from an OR transgene (*5×21-TAAR4Tg*) that includes a gene choice-promoting enhancer^24^ (Figure 6a). Several independent mouse lines showed over-expression of TAAR4 throughout the olfactory epithelium, and supernumerary glomeruli in the olfactory bulb (Figure 6a). Compared with *TAAR4-RFP* knockin mice, the highest expressing transgenic line had a 12-fold increase in OSN number and a 7-fold increase in the number of glomeruli (T4-RFP: 2.5±0.5 from reference [14] vs *5×21-TAAR4Tg*: 18.5 ± 1.2 (mean ± SD) glomeruli per bulb (n=8 bulbs). Importantly, individual OSNs that express TAAR4 from the *5×21-TAAR4* transgene have the same sensitivity as *TAAR4-RFP* OSNs, which express the endogenous TAAR4 locus (p=0.841, F=0.041; Sum-of-squares F-test; Figure 6b).

**Figure 6.**
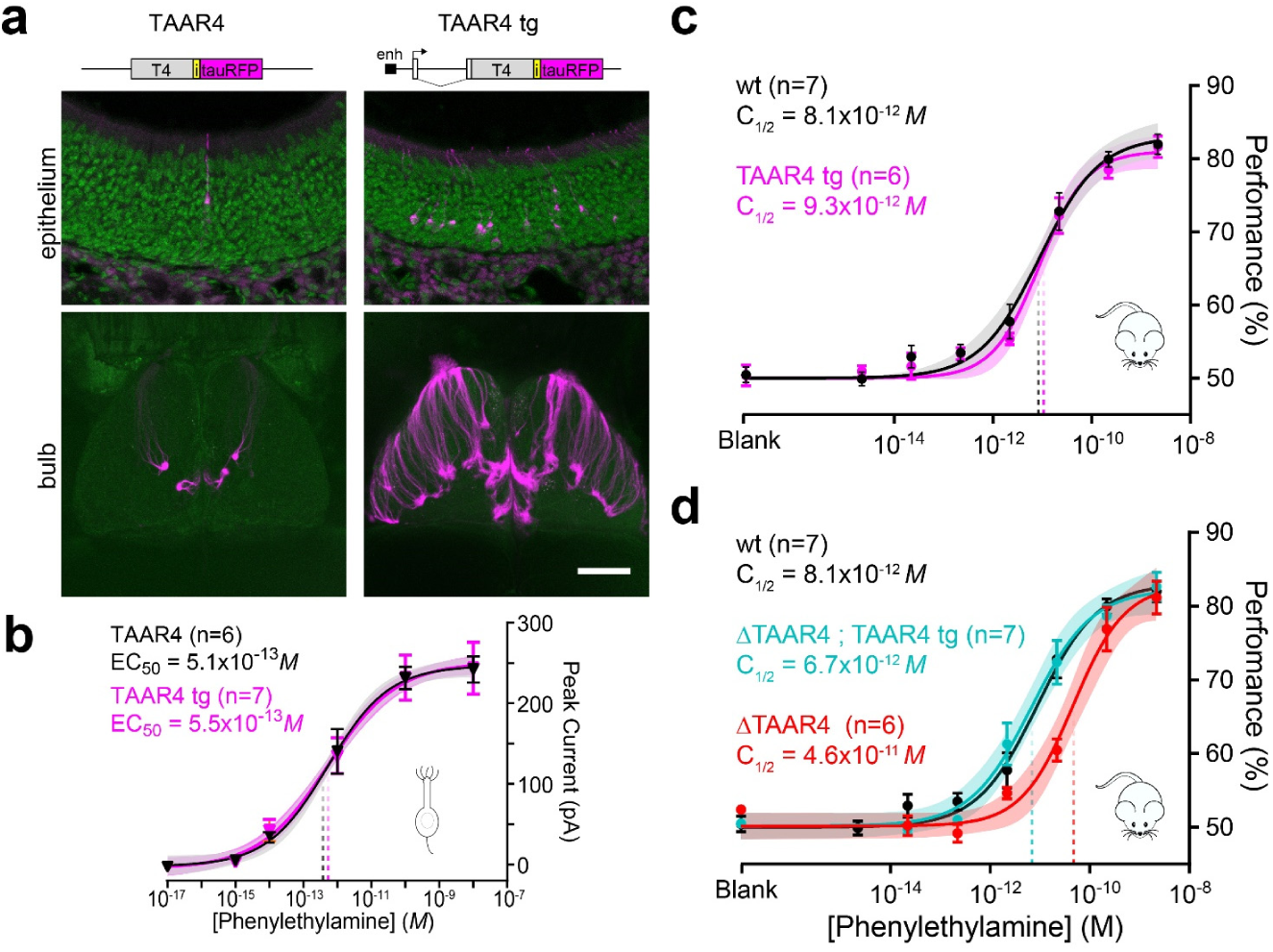
Over-expression of a threshold-determining TAAR does not enhance behavioral sensitivity. (**a**) Top: summary of transgenic strategy showing control gene targeted TAAR4 locus and transgene. The M71 transgene was modified by inserting a 5x homeodomain enhancer (enh) upstream of the transcription start site, and by replacing the M71 coding sequence with the TAAR4 coding sequence, an internal ribosome entry site (i) and a fluorophore (tauRFP). Bottom: Sections of the olfactory epithelium, and wholemounts of the olfactory bulb (dorsal view) in TAAR4-RFP and transgenic overexpressing mice. Scale bar is 50 μm in upper panels and 800 μm in lower panels. **(b)** Dose-response curves of OSN responses to phenylethylamine. Peak inward current was measured from single labeled neurons from TAAR4-RFP (black) and TAAR4 transgenic (magenta) mice. T4-RFP EC_50_ = 5.1 × 10^−13^ *M* (95% CI = 3.0-8.3 × 10^−13^*M*). *5×21- TAAR4Tg* EC_50_ = 5.5 × 10^−13^ *M* (95% CI = 2.5-11.6 × 10^−13^ *M*). **(c)** No effect of transgene expression on phenylethylamine sensitivity. Psychometric curves showing behavioral sensitivity (C_1/2_) of TAAR4 over-expressing transgenic mice (magenta), and wild-type littermates (black) to phenylethylamine. *wt* C_1/2_ = 8.1 × 10^−12^ *M* (95% CI: 4.6- 16.8 × 10^−12^ *M*), *5×21-TAAR4Tg* C_1/2_ = 9.3 × 10^−12^ *M* (95% CI: 6.0-15.9 × 10^−12^ *M*). **(d)** Effective transgenic rescue. Psychometric curves showing C_1/2_ of ΔTAAR4 mice (red) and ΔTAAR4 mice with the TAAR4 transgene (cyan) littermates. Wild-type (black) pyschometric curve is replotted from panel (c). *ΔT4* C_½_ = 4.6 × 10^−11^ *M* (95% CI = 2.4- 13.0 × 10^−11^*M*), *ΔT4*; *5×21-TAAR4Tg* C_½_ = 6.7 × 10^−12^ *M* (95% CI = 3.3-15.3 × 10^−12^*M*). Plots show mean +/- SE with a shaded 95% confidence interval.

Interestingly, TAAR4 overexpressing mice did not differ in their ability to detect phenylethylamine as compared to wild-type littermates (Figure 6c): C_1/2_ = 8.1 × 10^−12^ *M* for wild-type vs. 9.3 × 10^−12^ *M* for *5×21-TAAR4Tg* (p=0.710, F=0.139, Sum-of-squares F-test). This lack of effect is not attributable to a functional deficit of the transgene, since the transgene by itself was able to rescue the decrease in sensitivity caused by TAAR4 deletion (Figure 6d): C_½_ = 4.6 × 10^−11^ *M* for *ΔT4* vs. 6.7 × 10^−12^ *M* for *ΔT4*; *5×21-TAAR4Tg* rescue mice (p=0.001, F=11.01, Sum-of-squares F-test). These results argue that the normal complement of TAAR4 OSNs and glomeruli (typically 2 per olfactory bulb) does not limit behavioral sensitivity.

## Discussion

We have characterized the contribution of single olfactory receptors to behavioral detection in mice. Using TAARs as a model, we have defined the relative sensitivity of specific receptors to amines *in vivo*, and have precisely quantified behavioral detection deficits following the genetic deletion of multiple or even a single olfactory receptor. These experiments reveal that detection thresholds in mice are set solely by the single most sensitive olfactory receptor, and that sensitivity is not limited by typical numbers of sensory neurons and glomeruli allotted to each receptor.

Odor representations in the main olfactory system are thought to be highly redundant, with each OR making a small contribution to the representation of a given odor^25, 26^. It has been challenging to assess the contribution of individual ORs to odor detection, particularly in mammals. Studies in invertebrates have shed light on these issues^10, 11, 27, 28, 29, 30^. Deletion of single olfactory receptors in fly larvae affects chemotaxis^27, 28, 29^ and detection of a given odorant can be affected by highest affinity, as well as by lower affinity, receptors^10^. This suggests that detection threshold can be set by the most sensitive receptor, but also by pooling information from several high-sensitivity inputs. In addition, the exclusive expression of the most sensitive receptor for a given odorant increases sensitivity, while the exclusive expression of the highest and second highest affinity receptors together produced an intermediate sensitivity, suggesting a complex interaction among ORs to define threshold^11^.

In mammals, human studies have shown that polymorphisms in single OR genes correlate with variability in odor perception^31, 32, 33^, although it is not known if these ORs are expressed in the humans tested. In mice, there is a single example of an OR that is necessary for normal sensitivity to an odorant—Olfr1440 for the odorant muscone^9^. However, basic questions such as whether detection thresholds are determined solely by the most sensitive receptor, and how sensitivity at the level of OSNs correlates with behavioral thresholds have not been clearly answered. Our approach is unique because it combines genetic deletion of well-characterized, high-affinity receptors with rigorous behavioral thresholding.

We find that deleting the highest affinity receptor alters behavioral threshold, and that threshold is solely dependent on the most sensitive receptor. Note, that we have not tested whether the most sensitive TAAR alone (in the absence of all other ORs can support the same sensitivity. Overall, our findings are reminiscent of a “lower envelope” model in which behavioral threshold is dictated solely by the most sensitive afferents, in contrast to a pooling model in which behavioral threshold is determined by combining inputs of variable sensitivity^22^. The concept of pooling was originally put forth to address how reliable behavioral responses emerge from inherently noisy and unreliable single-neurons. It makes sense that a lower envelope scheme (across glomeruli) could work in olfaction, since the system may have solved the issue of noise in single sensory neurons by pooling functionally similar inputs via glomerular convergence. Our data indicate that information coming in via the most sensitive input (glomeruli) is sufficiently reliable to impart a given detection threshold. These findings further support the feasibility of models that emphasize the importance of the most sensitive receptors for olfactory coding^34, 35^.

Deletion of all olfactory TAARs caused an ~18-fold decrease in sensitivity to amines, rather than a complete loss of sensitivity. Our data indicate that the remaining receptors (ORs) may have sensitivities to particular amines that are within an order of magnitude of the TAARs. In fact, canonical ORs respond to amines^36^, albeit at much higher concentrations than our residual thresholds. Nevertheless, the fact that removing the entire TAAR repertoire does not produce amine anosmia underscores the functional redundancy of olfactory sensory inputs.

Our data also argue that a similar degree of functional redundancy can be seen in TAARs and ORs. The single TAAR deletions produced ~3 to 7-fold shifts in sensitivity to their preferred ligands, while the deletion of the class II OR (*Olfr1440*) produced a similar ~3-fold shift to muscone. Thus, single TAARs and ORs contribute similarly to sensitivity. While TAARs exhibit ultra-high sensitivity to amines ^12^, it is important to note that behavioral thresholds to the amines are within 10-fold of the those measured for methyl valerate and muscone, implying that ORs have comparable sensitivities as TAARs to their preferred ligands.

Our data confirm the observation that Olfr1440 sets the detection thresholds of mice to muscone, as previously described^9^. However, the sensitivity shift to muscone was reported to be several orders of magnitude—much larger than what we observe. The difference in effect size is like attributable to the different methods used to measure sensitivity. The previous study used an investigation-based assay in untrained, freely moving mice. This approach does not permit rigorous control of odor concentration, is difficult to quantify, and investigation times are confounded by factors not related to sensitivity (e.g. perceived intensity, valence, and motivation). In fact, assessing sensitivity of our TAAR cluster deletion mice using a similar investigation-based assay yielded an apparent shift in sensitivity of several orders of magnitude—much larger than what we observed via thresholding.

In our view, training-based thresholding provides a more accurate and relevant measure of sensitivity. In this approach, mice are well-trained and highly motivated to perform the detection task; it is known that performance in detection tasks improves with training and motivation^37, 38^. Furthermore, mice are tested in hundreds of trials for each odor concentration, providing a robust estimate of sensitivity despite inevitable behavioral variability. Our data provide a reliable measure of the detection limits of the olfactory system following specific genetic modifications. It is possible that under natural conditions (with untrained animals in a noisy olfactory environment) the shifts in sensitivity that we observe might manifest as marked deficits in odor localization.

The sensitivity of single TAAR glomeruli as measured by calcium imaging closely matched behavioral sensitivity as measured via thresholding. The indicator GCaMP3 exhibits robust fluorescence changes with neuronal activity, but may not reliably report single action potentials in vivo^39^. Nevertheless, under our imaging conditions, the convergence of thousands of olfactory sensory neurons to single glomeruli may facilitate visualization of fluorescence changes to near threshold odor concentrations. The concordance between these two measures is consistent with our observation that the highest affinity receptor sets the behavioral detection for the animal.

In addition, the shift in phenylethylamine threshold with TAAR4 deletion (~7-fold) closely matched the difference in sensitivity between TAAR4 and TAAR3 glomeruli measured via imaging. This observation argues that, in the absence of the highest affinity receptor (TAAR4), behavioral sensitivity is set by the second-highest affinity receptor (TAAR3). Thus, the concordance between the imaging and behavior is also consistent with the idea that behavioral sensitivity is set solely by the most sensitive receptor.

In mammals, the TAARs make up roughly 1% of the main olfactory receptor repertoire, yet they are highly conserved across species, perhaps reflecting a critical, universal role in olfactory function. The TAARs have been shown to mediate the detection of social cues or kairomones in mice, but such species-specific roles do not explain their broad conservation. Here, we show for the first time that the TAARs set the behavioral detection thresholds of mice to amines. Our data suggest that the TAARs may be evolutionarily conserved across vertebrates because they are the highest affinity amine receptors in the olfactory system.

Previous studies in mice show that deletion of TAAR genes abolishes apparently innate responses to amines—aversion or attraction^17, 19, 21^. One potential explanation for this loss of response is that TAAR deletion mice no longer detect the amines. However, our current study shows that TAAR deletion mice can still detect amines, albeit with reduced sensitivity. Unfortunately, a direct comparison of the stimulus concentrations in the present and earlier studies is not straight forward since the method of odor delivery is different (flow dilution olfactometry vs liquid-diluted odorants presented in open dishes). However, it is worth noting that ΔTAAR4 mice initially fail to avoid concentrations of phenylethylamine (presented in open dishes) that they subsequently detect and avoid following aversive conditioning (our unpublished observations). Thus, our data do not preclude the idea that TAARs play a crucial role in defining the identity, perceived intensity, and/or valence of amines, while also setting the detection threshold to these odorants.

Previous work has suggested that behavioral sensitivity may be determined in part by the overall number and convergence of OSN inputs^40, 41^. This assertion has been difficult to test, and related experiments have yielded conflicting results. Correlative studies in a number of species have noted a parallel between increasing OSN number and higher sensitivity^42, 43^. Likewise, experimental over-expression of two different ORs in mice (*Olfr151* or *OR1A1*) increased sensitivity as measured using an aversion-based, two bottle preference task^24^. On the other hand, a more pronounced over-expression (>95% of all OSNs) of an OR (*Olfr2*) yielded mice that did not differ in their investigation time to the corresponding ligand, octanal^44^. Interestingly, a similar over-expression (>95% of all OSNs) of a different OR (*M71*; *Olfr151*) produced mice that could not discriminate acetophenone from mineral oil, as measured in a Go/No-Go assay^45^^, 46^.

Our study is unique in that we have overexpressed a receptor (TAAR4) that is solely responsible for setting behavioral sensitivity. Despite this, overexpression of TAAR4 (in number of OSNs and glomeruli) did not change behavioral sensitivity. This result argues that a typical complement of receptor-defined OSNs and glomeruli does not limit detection. Our findings using odorant stimulation underscore the importance of previous work showing that mice can perceive the optogenetic stimulation of a single glomerulus^47^.

Convergence ratio (OSN to mitral/tufted cell) is a key factor in models that propose increases in sensitivity with OSN number^40^. However, modulating convergence ratios remains an experimental challenge, and it is possible that increasing convergence might shift sensitivity. In addition, it should be noted again, that our results are collected using a highly-controlled odor source and well-trained mice that perform thousands of trials. Thus, it is possible that receptor over-expression might increase sensitivity under more natural conditions (untrained animals that encounter fluctuating olfactory stimuli on the presence of background odors). Regardless, our data provide evidence that, in a well-controlled setting, a typical complement of highest-affinity glomeruli is sufficient to translate the sensitivity of specific molecular receptors into a behavioral detection threshold.

Overall, our results argue that, among the >1,000 main olfactory GPCRs, the TAARs are the most sensitive to amines. Thus, it seems likely that this phylogenetically distinct class of aminergic receptors was co-opted to an olfactory function for this high sensitivity. The importance of TAARs for amine detection may explain their evolutionary conservation. More broadly, the fact that a single olfactory receptor gene (either a TAAR or an OR) can make a significant contribution to behavioral sensitivity suggests a mechanism by which olfactory receptors could be under selective pressure, thus shedding light on how vertebrates maintain large olfactory receptor repertoires over evolutionary time.

## Methods

All procedures involving animals were approved by the Northwestern University Institutional Animal Care and Use Committee.

### Mouse Strains

The generation of *ΔTAAR4-Venus*, *TAAR4-IRES-tauCherry*, *TAAR3-IRES-tauVenus* and the TAAR cluster deletion allele (*ΔT2-9*) were previously described^14^. The *ΔTAAR5-LacZ* deletion allele was obtained from KOMP (*Taar5^tm1(KOMP)Vlcg^*) and the selection cassette was removed in the germline by Cre-mediated recombination after crossing with *HPRT-Cre*/129 mice.

The targeted mutation *ΔTAAR3-GapTeal* was generated by replacing the receptor coding sequence in the TAAR3 targeting vector^14^ from the start codon to the stop codon with a fragment encoding the teal fluorescent protein, mTFP1^48^ fused to the N-terminal 20 amino acids (palmitoylation signal) of zebrafish GAP43^49^. The self-excising neomycin selection cassette ACNF was inserted into the AscI site to generate the final targeting vector. The targeted mutation *OMP-GCaMP3* was generated by cloning the coding sequence of GCaMP3^39^ flanked by AscI sites into the targeting vector for the olfactory marker protein (*omp*) locus^50^ so that the coding sequence of OMP is replaced by that of GCaMP3. The self-excising neomycin selection cassette *ACNF* was inserted 3’ of the coding sequence.

All targeting vectors were electroporated into a 129 ES line, and clones were screened for recombination by long range PCR. Chimeras were generated from recombinant clones using C57BL/6J host embryos. All gene targeted mouse strains are on a mixed B6/129 background. The *OMP-GCaMP3* strain is available from The Jackson laboratory (Stock #029581).

*5×21-TAAR4Tg* : A transgene consisting of a 6.8 kb fragment of the M71 locus was modified by 1) insertion of a 5x homeodomain sequence ^24^ placed 490 bp upstream of the transcription start site, and 2) by replacing the *M71* coding sequence with the coding sequence for mouse TAAR4 flanked by Asc I sites and preceded by a Kozak consensus sequence (*GGCGC*<u>*GCC*ACCATG</u>). The coding sequence was followed by an internal ribosome entry site and tauCherry (a fusion of bovine tau and monomeric cherry fluorescent protein^51^. The transgene was column purified and injected into C57BL/6J zygotes to obtain transgenic founders. Experimental animals were maintained on a C57BL/6J background.

*ΔOlfr1440*: A guide RNA was designed to target a PAM sequence in the 5’ coding sequence of the *olfr1440* gene, and the recognition sequence cloned into the pX458 (Addgene #48138). The gRNA along with Cas9 RNA was injected into C57BL/6 zygotes. Founders were screened by PCR and sequencing. A modified allele was isolated with a 71 bp deletion starting at nt 44 of the *olfr1440* coding sequence. This produced a frameshift after the first 14 amino acid residues, encoding a predicted 26 amino acid peptide (MPGGRNSTVITKFIHSGVELGPHHLD-STOP). The allele was backcrossed for three generations on a C57BL/6J background, then intercrossed to produce experimental animals.

### Head-Fixed Odor Thresholding

Head-bars were surgically implanted on all experimental animals. Prior to surgery, wild-type and mutant littermates were housed in same sex groups until 10-14 weeks of age. Mice were anesthetized with isoflurane (2-3%) in oxygen and administered buprenorphine (0.1 mg/kg) as analgesic, and bupivacaine (2 mg/kg) as a local anesthetic at the incision site. The animals were secured in a stereotaxic head holder (Kopf). Two micro-screws were placed in the skull, and a custom-built titanium head-bar (<1g) was attached to the skull and screws using Vetbond cyanoacrylate glue and cemented in place using dental cement (Dental Cement, Pearson Dental Supply).

After surgery, mice were individually housed and given at least two days for recovery before water restriction. Water restricted mice (> 2 weeks at 1 ml of water per day) were trained to report the detection of odor in a Go/No-Go task in a custom built apparatus. In each cohort of mice, equal numbers of mutant and wild-type littermates were run without knowledge of genotype. Mice were placed in a custom holder with their noses 1 cm from an odor port. The odor port was mounted on a concave base which housed the lick tube and vacuum connection to remove excess odor. Licks were detected electronically using a custom-built lick circuit. Water delivery was controlled by a solenoid valve connected to a small water reservoir. The animal holder and odor port were mounted on a breadboard inside an 18×18×18” custom-made sound proof box.

Odorants were diluted in water (amines/methyl valerate) or mineral oil (muscone) and delivered using a flow dilution olfactometer^20^. The saturated headspace of each diluted odorant was further diluted 10-fold via flow prior to reaching the animal. The maximal concentration of each odor tested is as follows: isopentylamine: 4.8×10^−6^ *M* (2% liquid dilution); phenylethylamine: 4.4×10^−8^ *M* (2% liquid dilution); trimethylamine: 4.9×10^−5^ *M* (2% liquid dilution); methyl valerate: 6.1×10^−7^ *M (*1% liquid dilution); and muscone: 2.5×10^−9^ *M* (undiluted). A dual synchronous three-way solenoid valve connected the olfactometer and the purified air lines (with the same flow rate) to an exhaust line and the odor port. Care was taken to ensure that both lines were impedance matched to prevent pressure spikes during odor delivery. During stimulus application, the dual-synchronous valve swapped the flow to the animal from clean air to balanced, diluted odorant. Disposable 40 ml amber glass vials filled with 5 ml of the odorant dilution were attached to the olfactometer manifolds and pressurized before the start of the first trial. Mice were tested only once per day with a single odor concentration (to limit contamination) and were thresholded to a maximum of two odors total (to prevent over-training and solving the task using non-odor cues – see below). A custom Arduino-based behavioral controller coordinated the trial structure and monitored licking. A custom Matlab script sent trial parameters to the behavioral controller and olfactometer.

Behavioral training consisted of two stages. Stage 1 – the mice were trained to receive a water reward if they licked during the 2 second stimulus period (signaled by an LED). The inter-trial interval was steadily increased from 1.5 to 5 seconds over the course of several sessions. Mice were exposed to a constant stream of clean air (1 L/min) during this training. Stage 2 – mice were trained in a Go No-Go odor detection task. A blank olfactometer vial (5 ml of Milli-Q water) served as the Go stimulus while a vial containing a high concentration of the target odor (100 µL of odor in 5 mL of Milli-Q water) served as the No-Go stimulus. Correct responses during the 2 second stimulus window were rewarded with water (1.5-2 µL) and/or a short inter-trial interval (5-9 seconds). Incorrect responses were punished with a long inter-trial interval (15-19 seconds). To minimize early false alarms, the first ten trials were Go trials and the session was terminated after the mice missed three Go trials in a row. Sessions typically lasted 250-600 trials. Behavioral performance was determined by the number of correct responses (hits+correct rejections) divided by the total number of trials (after the initial Go trials). Mice learned this task quickly and usually performed above 90% on the second session. Upon reaching criteria (two sessions above 90% correct), mice were subsequently thresholded. A given cohort was used to collect a full threshold series for at most two different odorants.

For thresholding, the olfactometer was loaded with three blank (Go) vials, three diluted odor (No-Go) vials, and a single blank (No-Go) vial. Each vial was replaced daily and their positions were randomized. This blank No-Go vial served to test whether the mice were using cues other than the presence or absence of the target odor to maximize performance. Performance enhanced by non-odor cues was determined by the ability of the mouse to correctly reject the blank No-Go vial at a frequency higher than the percentage of misses (not licking during a Go stimulus). If this occurred, the session was excluded from the analysis. If this occurred three times over the course of an experiment, the mouse was removed from the experimental group. Since this check is included in our analysis, the maximum performance a mouse can attain using only odor cues is approximately 85%. At the end of each day, the olfactometers were flushed with 70% isopropanol and dried with pressurized clean air overnight.

Several additional modifications were necessary to threshold mice to trimethylamine due its high vapor pressure. To minimize fluctuations in odor concentration, the stimulus period was shortened to 1 second, a fourth odor vial was included, and diluted trimethylamine was replaced after each mouse. To minimize contamination, the inter-trial intervals were increased to 12-17 seconds for a correct response and 22-27 seconds for an incorrect response. In addition, high pressurized clean air (15 psi) was forced through the olfactometer during each inter-trial interval to remove any residual odorant.

Behavioral performance for each odor and genotype was fitted to a Hill function

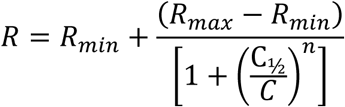

where *R* is the behavioral accuracy, *C* is odor concentration, *C_½_* is the concentration at half-maximal performance, and *n* is the Hill coefficient. The C_½_ values were compared between genotypes using a sumof-squares F-test (Prism Graphpad).

### Awake *In Vivo* Imaging

Imaging was performed in awake, 3-6 month old mice that were heterozygous for the *OMP-GCaMP3* allele and compound heterozygous for the *TAAR4-RFP* and *TAAR3-YFP* alleles. Prior to imaging, mice were implanted with a head bar (as describe above) and a chronic imaging window. Mice were administered dexamethasone (2 mg/kg) to reduce cerebral edema, and the bones overlying the olfactory bulbs were thinned to transparency using a dental burr. Black Ortho Jet dental acrylic (Lang Dental Manufacturing) was used to secure the head bar and to form a chamber over the imaging area. The bone overlying the olfactory bulbs was covered with multiple thin layers of prism clear cyanoacrylate glue (Loctite #411) as described^50^. Following complete recovery from surgery, mice were placed on a water restriction schedule (1 ml/day). After seven to ten days of restriction, mice were slowly habituated to the imaging setup where they were trained to lick for a water reward. During imaging sessions, mice were secured via the head-bar in a custom-built animal holder and running wheel.

Glomerular imaging was performed using wide field epifluorescence microscopy, and GCaMP signals recorded using a CCD camera (NeuroCCD SM256; RedShirtImaging) at 25 Hz with a 4x temporal binning. Light excitation was provided using a 200 W metal-halide lamp (Prior Scientific) filtered through standard filters sets for RFP (Chroma 49008), GFP (Nikon 96343), and YFP (Chroma 86001). Each recording trial was 16 s consisting of a 6 s prestimulus interval, a 4 s odor pulse and a 6 s poststimulus interval.

Odorants were diluted in water, and then further diluted and delivered using a computer-controlled, flow dilution olfactometer^20^ as described above, but modified to present odor from a single interchangeable vial. This modification was necessary to prevent contamination of the olfactometer by high concentrations of a given odorant. A single disposable 40 ml amber glass vials (Thermo Scientific) filled with 5 ml of the odorant dilution was attached directly to one mass-flow controller of the olfactometer. This odorized air was further diluted using a second mass-flow controller and a Teflon mixing chamber. A dual synchronous three-way solenoid valve (Neptune Research, SH360T042) presented either odorized air from the Teflon mixing chamber or clean air (with the same flow rate) to the animal’s nose via a custom-built odor port. An exhaust line of the same resistance limited pressure spikes and allowed the odor to stabilize before the beginning of each trial. Only one odorant was tested per day to avoid cross contamination, and odor trials were interleaved with clean air trials to identify potential contamination. If contamination was observed, the mixing chamber and solenoid valve were replaced and the experiment continued.

Response maps were obtained by subtracting a 1.6 s temporal average preceding the stimulus from a 0.5 s temporal average at the maximum response during the first 2 seconds of the stimulus. Images were processed and analyzed in Neuroplex (RedShirtImaging) and Image J ^52^ software. Response amplitudes of TAARs 3, 4, and 5 were measured from regions of interest drawn around each glomerulus. Responses for each odor were fitted to a Hill function

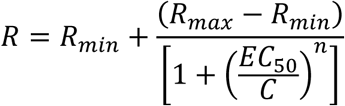

where *R* is the response amplitude, *C* is odor concentration, *EC_50_* is the concentration at half-maximal response, and *n* is the Hill coefficient. The EC_50_ values were compared between receptors using a sum-of-squares F-test (Prism Graphpad).

### DREAM Assay

Male C57BL/6J mice (40-60 days old) were individually housed 24 hours prior to the experiment and provided with food and water *ad libitum*. On the day of the experiment, mice receiving odorant exposure were placed in new clean cages connected to the olfactometer. We adapted our custom built flow dilution olfactometer to deliver odor sequentially to 5 individual mouse cages for 1 minute each for 24 hours. Excess odor was exhausted passively. To minimize odor loss, the olfactometer was loaded with 10 vials (each with 5 ml of diluted odorant). Control mice were placed in new clean cages in an adjacent odor free room. After 24 hours, both control and odorant exposed mice were euthanized and the olfactory epithelium was immediately collected for RNA extraction using the RNAeasy kit (Qiagen). Isolation was performed according to the manufacturer’s protocol RNA was treated with Ambion DNase I kit (Life Technologies) to eliminate genomic (DNA) contamination. The quality and concentration of the RNA was determined (2100 Bioanalyzer; Agilent Technologies) prior to cDNA synthesis (Invitrogen SuperScript^TM^ III Reverse Transcriptase; 300 ng of total RNA). Transcript expression was measured using real-time qPCR (Bio-Rad CFX384) using the iQ^TM^ SYBR^®^ Green Supermix (Bio-Rad). Gene-specific primers were designed and validated for efficiency. The homology among TAAR 8a, 8b, and 8c was too high to permit the design of specific primers. Cycling parameters were: 95°C for 3 minutes, followed by 40 cycles of 95°C for 15s and 59°C for 45s. All reactions were run in triplicate. Raw Ct values for each TAAR were normalized to the geometric mean of the broadly expressed reference gene *gurb* and the olfactory tissue-specific gene *gnal*. Normalized control and odor exposed Ct values were compared using a General Linear Mixed Model with pairwise comparisons.

### Histological preparations

For the preparation of the olfactory epithelium sections, *TAAR4-RFP* and *5×21-TAAR4Tg* mice (20-30 days old) were anesthetized and transcardially perfused with heparinized saline followed by 4% paraformaldehyde. The rostrum was dissected and postfixed at 4°C for 1 hour followed by 0.5M EDTA decalcification and 30% sucrose cryoprotection (both overnight at 4°C). OCT-embedded noses were sectioned at 16µm. For both strains, all the TAAR4-expressing OSNs were counted on representative sections using confocal microscopy.

For the wholemount analysis of the number of glomeruli in the *5×21-TAAR4Tg* strain, the genetically encoded fluorescent marker (RFP) was visualized without fixation by confocal microscopy. Olfactory bulbs and brain were removed from the skull, embedded in low-melting temperature agarose, and all surfaces of the olfactory bulb were scanned using a Leica SP8 confocal microscope. A detailed analysis of the number of glomeruli in the *TAAR4-RFP* strain was previously published^14^.

### Statistics

No statistical methods were used to predetermine sample sizes, but sample sizes were determined based on numbers of animals required to obtain statistically significant effects in pilot experiments. Data collection and analysis were performed blind to the conditions of the experiments. Statistical tests were performed using Prism (GraphPad) or SPSS (IBM) and are indicated in the figure legends. All statistical tests were two-tailed. Assumptions about normal distributions were tested using the Shapiro-Wilk normality test, and assumptions about equality of variances were tested using an F-test. A Generalized Estimating Equations Model was used to fit a model to repeated categorical responses and to test for differences between groups.

## Acknowledgements

This work was supported by grants from the NIH/NIDCD and DARPA. We thank Matt Smear, Roman Shusterman and Yevgeniy Sirotin for their input and technical help during early phases of the project, Nao Uchida for discussions regarding sensory thresholds, Admir Resulaj for technical help with olfactometry, Sarah Kaye, Dillon Cawley, Hardik Patel, Tiffany Teng, and Amanda Menzie for technical support, and William Kath and Rosemary Braun for statistical advice. We acknowledge Lynn Doglio and the Transgenesis and Targeted Mutagenesis Laboratory at Northwestern University for transgenic mouse production.

